# Multiscale structural anisotropy in pulvinus motor organs of the sensitive plant *Mimosa pudica*

**DOI:** 10.1101/2022.02.28.482281

**Authors:** David A. Sleboda, Anja Geitmann, Reza Sharif-Naeini

## Abstract

Pulvini are joint-like motor organs that power active leaf movement in many plants. Multiple structural specializations spanning subcellular, cellular, and tissue scales of pulvinus organization have been described; however, the impacts of multiscale mechanics on pulvinus physiology remain poorly understood. To investigate the influence of multiscale morphology on turgor-induced deformation, we visualized *Mimosa pudica* pulvinus morphology at multiple hierarchical scales of organization and used osmotic perturbations to experimentally swell pulvini in incremental states of dissection. We observed directional cellulose microfibril reinforcement, oblong, spindle-shaped primary pit fields, and flattened, disk-like cell geometries in the parenchyma of *M. pudica*. Consistent with these observations, isolated parenchyma tissues displayed highly anisotropic swelling behaviors indicating a high degree of mechanical anisotropy. Swelling behaviors at higher scales of pulvinus organization were also influenced by the presence of the pulvinus epidermis, which displayed oblong epidermal cells oriented transverse to the pulvinus long axis. Our findings indicate that structural specializations spread across multiple hierarchical scales of organization guide hydraulic deformation of pulvini, suggesting that multiscale mechanics are crucial to the translation of cell-level turgor variations into organ-scale pulvinus motion *in vivo*.

## Introduction

Pulvini are fleshy swellings located at the bases of petioles, leaves, and leaflets in many plants. They are sites of growth-independent motion and facilitate active, reversible bending of leafy plant appendages. Anatomically, pulvini are roughly cylindrical in shape and consist of a flexible core of vasculature surrounded by a thick cortex of parenchyma cells. During active deformation, bulk migration of water occurs between extensor and flexor parenchyma cell groups located on opposing sides of the central vasculature core (Tamiya et al., 1988), resulting in a length differential between the two groups that induces long-axis pulvinus bending. Pulvinus-driven leaf folding can occur gradually over the course of hours under the influence of a circadian rhythm, or rapidly as a thigmonastic response to touch, shaking, or wounding of a plant. Active pulvinus bending is associated with efflux of K^+^ and Cl^-^ ions from parenchyma cells (Kumon and Suda, 1984), offloading of sugar from vascular elements (Fromm and Eschrich, 1988a), and consumption of ATP (Fromm and Eschrich, 1988b), indicating that pulvinus motion is associated with active transport of ions and other osmolites to which water is attracted (Scorza and Dornelas, 2011).

While multiple theoretical models of pulvinus physiology provide insight into the biochemical mechanisms that power active movement of fluid between parenchyma cell groups (e.g. Kagawa and Saito, 2000; Morillon et al., 2001; Kwan et al., 2013; Wang and Li, 2020), relatively little is known regarding the mechanisms by which volumetric changes at the cellular level are translated into physiologically useful motion at the organ scale. Multiple morphological features have been identified which have the potential to bridge this gap. In pulvini of the scarlet runner bean *Phaseolus coccineus L*., observations with polarized light microscopy suggest that the walls of parenchyma cells are reinforced by cellulose microfibrils oriented primarily perpendicular to the long axis of the pulvinus, confining turgor-induced cell deformation to the long axis of the organ where its effect on bending is greatest (Mayer et al., 1985). In pulvini of *Phaseolus coccineus L*. and of the common bean *Phaseolus vulgaris*, parenchyma cells are flattened in the direction parallel to the long axis of the pulvinus, displaying approximately disc-shaped morphologies that are also conducive to deformation preferentially along the long axis of the organ (Mayer et al., 1985; Koller and Zamski, 2002). In pulvini of *Albizia julibrissin, Phaseolus vulgaris*, and the sensitive plant *Mimosa pudica*, wrinkles in the pulvinus surface run preferentially transverse to the organ long axis and have been identified as structural features that facilitate longitudinal deformation of the pulvinus epidermis without sacrificing circumferential stiffness (Satter et al., 1970; Koller and Zamski, 2002; Song et al., 2014; Mano and Hasebe, 2021). These structural features have the potential to physically guide hydraulic deformation of pulvini, ensuring that volumetric changes at the cellular level are ultimately translated into long-axis tissue deformations that induce macroscopic organ bending. Despite the functional importance of this role, a multiscale study surveying structural anisotropy across scales of pulvinus organization within a single plant species has not been carried out, and the distinct contributions of cell wall ultrastructure, cell geometry, and epidermis morphology to the management of hydraulic deformation remain unclear.

Here we disentangle the influences of cell wall ultrastructure, cell geometry, and epidermis morphology on turgor-induced deformation in the pulvini of *Mimosa pudica*, a touch-sensitive plant which displays both rapid thigmonastic and slow circadian leaf movements. Using a combination of scanning electron and confocal microscopy, we visualized *M. pudica* pulvinus morphology at multiple hierarchical scales of organization and characterized the degree of structural anisotropy present at each. Additionally, we used osmotic bathing solutions to experimentally swell live, isolated *M. pudica* pulvinus tissues in various states of dissection. This approach allowed us to characterize the degree of anisotropic tissue deformation that results from purely isotropic loads of intracellular fluid pressure. Differences in swelling behaviors across scales of organization are discussed in relation to the anisotropic structural features observed.

## Materials and Methods

All pulvinus organs and tissue samples were harvested fresh from *Mimosa pudica* plants grown under laboratory conditions. Plants were grown from seed in a growth chamber with a 12-hour light-dark cycle at 21-23°C and 60% humidity. Plants were watered approximately once every 3 days and fertilized monthly.

### Epidermis morphology

The surface morphology of the pulvinus epidermis was visualized using a Hitachi TM-1000 scanning electron microscope (SEM). Primary, secondary, and tertiary pulvini located at the bases of petioles, leaves, and leaflets, respectively, were removed from plants and dehydrated in a graded series of ethanol. Isolated pulvini were then critical point dried in liquid CO2 and mounted in lateral view on aluminum stubs with carbon double-sided tape. Mounted organs were sputter coated with a 4 nm layer of gold and palladium and imaged at an excitation voltage of 15 kV. Aspect ratios and orientations of epidermal cells were measured manually from scanning electron micrographs of primary pulvini using ImageJ (Schindelin et al., 2012). The positions of epidermal cells along the lengths of pulvini were defined relative to their distance from the pulvinus base, measured along a path traversing the central axis of pulvinus and petiole (see yellow line depicted in fig. 2a). Total path length examined was determined by doubling the distance from the base of the pulvinus to the center of the knuckle-like protuberance immediately distal to the pulvinus (indicated as length 0.50 in fig. 2a), such that roughly equal lengths of pulvinus and petiole were examined. To facilitate comparison of pulvini of varying size, distances of epidermal cells from the pulvinus base were expressed as portions of the total path length examined for each organ. Epidermal cell orientation was measured relative to the orientation of the central path.

### Cell wall ultrastructure

The ultrastructures of individual parenchymal cell walls were visualized using an environmental scanning electron microscope (FEI Quanta 450 FE-ESEM, FEI Co.) at the McGill University Facility for Electron Microscopy Research. In order to observe the inner structures of individual cell walls, primary pulvini were decellularized using an adapted protocol for the generation of acellular, cellulosic tissue engineering scaffolds from plant material (Adamski et al., 2018). Fresh pulvini were fixed overnight in 4% formaldehyde in 0.1 M phosphate buffer, pH 7.2 (Kwan et al., 2013) and then bisected longitudinally using a fine razor blade under a stereomicroscope. Fixed, bisected pulvini were immersed in an aqueous solution of 3% NaOH, 5% bleach for approximately 20 hours at room temperature until they appeared uniformly white and translucent under transmitted light illumination, indicating maceration of cellular material. Pulvini were then gently rinsed in distilled water for 24 hours, dehydrated in 100% ethanol, critical point dried in liquid CO2, mounted on aluminum stubs with carbon double-sided tape, and sputter coated with a 4 nm layer of gold and palladium. Exposed internal faces of the bisected pulvini were imaged at an accelerating voltage of 10 kV, providing lateral views of the internal structure of the decellularized organs.

The spatial orientations of cellulose microfibrils composing parenchyma cell walls were measured from high-magnification (~50,000-65,000x) scanning electron micrographs. Cellulose microfibril orientation was quantified using the Directionality plugin in ImageJ (Liu, 1991). Analysis via either the Fourier components or local gradient modes of the plugin yielded similar results, and results from the local gradient method are reported here. Orientations of identifiable structures in each cell wall micrograph were expressed as a histogram with portion of structures observed on the Y axis and angular orientation in degrees on the X axis. The peak of a gaussian curve fit to the collective histogram compiled from all micrographs provided a quantitative metric of the predominant orientation of cellulose microfibrils across samples, and the standard deviation of the gaussian curve provided a metric of histogram width, i.e., the dispersion of microfibril orientations about the predominant angle.

Aspect ratios and spatial orientations of primary pit fields, which physically connect adjacent parenchyma cells, were also measured from scanning electron micrographs manually using ImageJ. Pit fields were identified as matted areas of cell wall perforated by numerous plasmodesmata. The spatial orientations of both pit fields and cellulose microfibrils were measured relative to the pulvinus long axis, as indicated by the orientations of nearby vasculature running longitudinally through the organ core.

### Multiscale osmotic swelling

To explore differences in osmotic swelling behaviors across scales of pulvinus organization, pulvini in incremental states of dissection were imaged in aqueous mannitol baths of varying tonicity. Mannitol is a biologically inert molecule and has been used previously to induce osmotic deformation of pulvinus tissues (Mayer et al., 1985). Bathing in relatively dilute, hypotonic mannitol solutions produces an osmotic gradient that favors movement of water into cells, resulting in tissue expansion, while bathing in relatively concentrated, hypertonic mannitol solutions favors movement of water out of cells, resulting in tissue shrinkage.

Three preparations of pulvinus organ were subjected to graded osmotic manipulation: cylindrical sections of pulvini, cylindrical sections skinned of their epidermis, and isolated blocks of parenchyma tissue. Cylindrical sections of pulvinus organ were produced by removing the proximal and distal ends of isolated pulvini using a fine razor blade under a stereomicroscope, resulting in sushi roll-like sections of pulvinus organ 1000-1500 μm long and 1000-1400 μm wide. Sections comprised a core of vasculature surrounded concentrically by a thick parenchyma cortex and thin enveloping epidermis. Removal of the epidermis from a subset of sections was accomplished via manual skinning with a fine razor under a stereomicroscope and produced cylindrical sections containing only the vasculature core and surrounding parenchyma cortex. Isolated parenchyma blocks were taken from the relatively large extensor (i.e. ventral or abaxial) cortexes of primary pulvini and were free of both vascular and epidermis components. Flexor (i.e. dorsal or adaxial) cortexes of *M. pudica* pulvini were found to be too small to reliably yield parenchyma samples via manual dissection under a stereomicroscope, and were not examined in the current study.

Dissected pulvinus preparations were transferred through a graded series of osmotic mannitol baths ranging in concentration from 0 to 878 milliosmoles (mOsm). Solution concentrations were calculated and prepared via serial dilution of measured quantities of mannitol dissolved in distilled water. Following an acclimation time of at least 30 minutes in each osmotic bath, samples were imaged in both lateral and ventral views using a digital camera (Zeiss AxioCam MRc, Carl Zeiss AG) mounted on a stereomicroscope (Zeiss SteREO Discovery V8, Carl Zeiss AG). Widths, lengths, and heights of dissected pulvinus preparations were measured at their maxima from stereomicrographs using ImageJ. Measurements of dissected pulvinus preparations were normalized to dimensions in the most hypertonic osmotic bath employed (878 mOsm). Statistically significant changes in tissue size were identified via two-tailed, one sample t-tests comparing normalized dimensions in the most hypertonic (878 mOsm) and most hypotonic (distilled water) bathing solutions.

### Parenchyma cell geometry

The geometries of live parenchyma cells were visualized using a laser scanning confocal microscope (LSM710, Carl Zeiss AG). Thin sections of fresh parenchyma tissue were collected from the relatively large extensor (i.e. ventral or abaxial) cortexes of primary pulvini using a pair of razor blades spaced by ~400 μm of double-sided tape. Thin sections were prepared for confocal microscopy by staining briefly (7-9 minutes) with 0.5 mg/ml propidium iodide (PI) in osmotically isotonic bathing solution. PI binds to cell wall pectin and is a common stain for live imaging of plant anatomical features (Bidhendi et al., 2020). Isotonic bathing tonicity was determined by bathing parenchyma tissue in a graded series of aqueous mannitol solutions and measuring changes in tissue size. Osmotic baths more concentrated than 384 mOsm of mannitol were found to cause shrinkage of freshly dissected parenchyma tissues, while osmotic baths less concentrated than 384 mOsm caused expansion. After fluorescent staining, parenchyma sections were rinsed briefly of unbound PI and then excited with a 543 nm Helium-Neon laser at 5% power. Fluorescence emission in a range of 554-711 nm was captured from parenchyma cells deep to the sectioned tissue surface. Cells with a fluorescing nucleus indicating penetration of PI into the cell protoplasm were considered damaged by sectioning and excluded from the study. Optical sections 2.4-6.4 μm thick were captured at the midlines of cells of interest, located by changing focus along the z-axis until the largest apparent cell dimensions were presented. Slice thickness was changed by manipulating the size of the confocal pinhole.

To examine the influence of increased intracellular fluid volume on the geometry of individual parenchyma cells, parenchyma tissues visualized via confocal microscopy were initially imaged in osmotically isotonic bathing solution and subsequently re-imaged in osmotically hypotonic baths of distilled water. Isotonic and hypotonic baths were exchanged without moving parenchyma tissue samples from the confocal stage, allowing visualization of individual parenchyma cells of interest before and after osmotic swelling. An acclimation time of a least 5 minutes in hypotonic solution was given prior to re-imaging of individual cells. The lengths, heights, widths, lateral perimeters, and frontal perimeters of individual parenchyma cells were measured from confocal micrographs in ImageJ and normalized to dimensions in osmotically isotonic baths. Changes between isotonic and hypotonic solutions were expressed as percentages, and significant changes in cell dimensions between the two bath conditions were identified as those significantly different from zero in a two-tailed one sample t-test. Lateral aspect ratios of parenchyma cells were calculated by dividing cell length (measured in the direction parallel to the long axis of the pulvinus) by cell height (measured along the dorsal-ventral axis of the organ). Frontal aspect ratios of cells were calculated by dividing cell width by cell height. Non-circular aspect ratios were defined as those that differed significantly from 1.0 in a two-tailed one sample t-test.

### Anatomical descriptors

All anatomical descriptors employed in the current study are defined relative to organ-scale pulvinus anatomy. Lengths are measured along the long axis of the pulvinus, defined by the orientation of the vasculature tissue running longitudinally through the organ center. Heights are measured along the dorsal-ventral axis of the pulvinus, perpendicular to the neutral axis of active pulvinus bending. Widths are measured laterally across the cross-section of pulvini. These terms are employed across all hierarchical scales of organization examined. For example, the length of an individual parenchyma cell refers to size measured along the long axis of the pulvinus it composes.

## Results

Pulvini of *Mimosa pudica* displayed structural anisotropy at all scales of organization examined. Epidermal cells bounding the surfaces of pulvini were oblong in shape with long axes oriented perpendicular to the organ long axis (Fig. 1). Wrinkles in the pulvinus surface were oriented predominantly transverse to the pulvinus long axis, i.e., parallel to the long axes of epidermal cells. Surface wrinkles were confined exclusively to pulvinus organs, and were not apparent on neighboring surfaces of stems, petioles, or leaflets.

**Figure 1:**
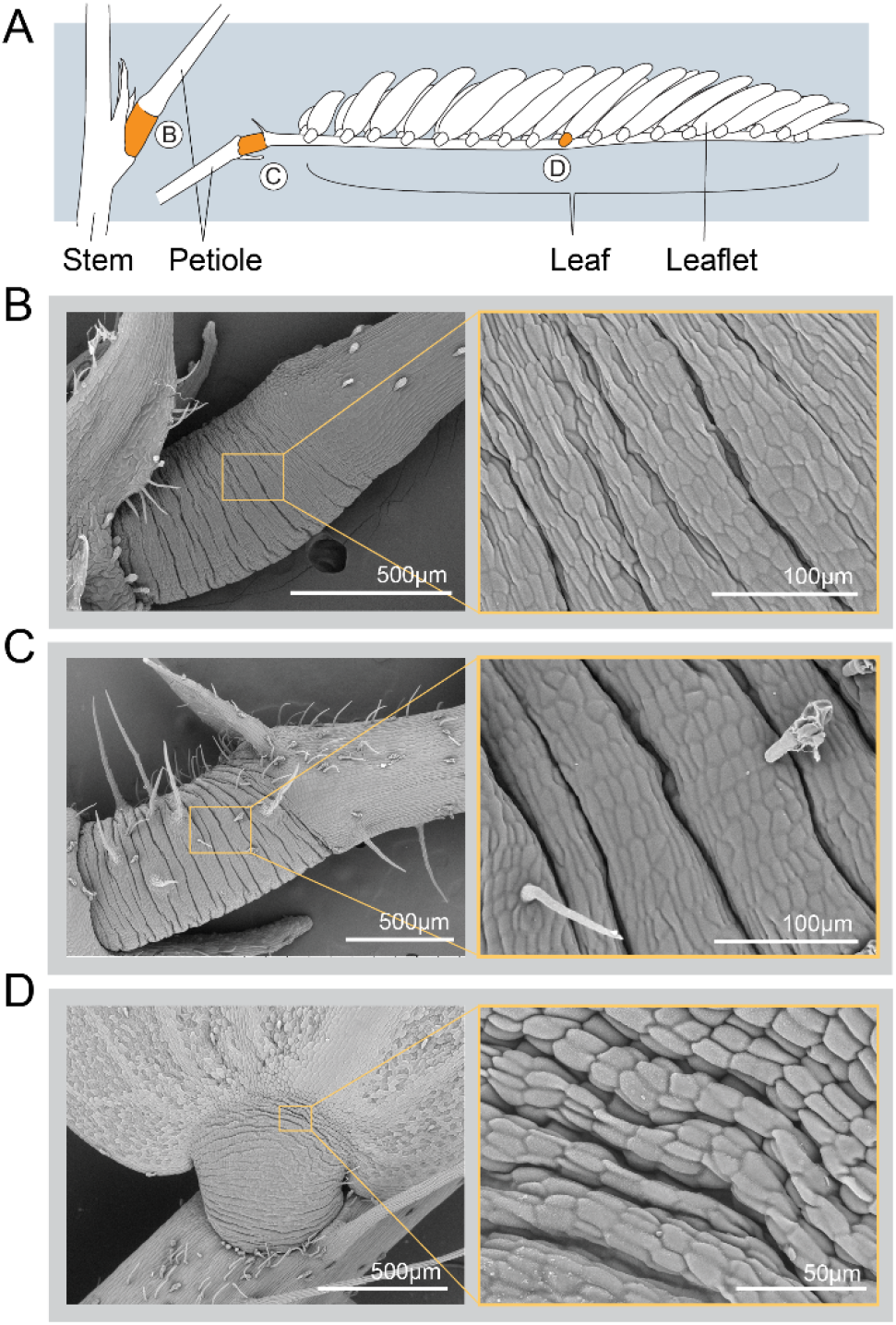
Pulvinus locations and external morphologies. [A] Line drawing depicting the locations of primary (B), secondary (C), and tertiary (D) pulvini of *Mimosa pudica*. [B,C,D] Scanning electron micrographs of pulvinus organs identified in A. Inset micrographs show epidermis morphology at higher magnifications. In the immediate regions of pulvini, epidermal cells are oblong and oriented perpendicular to the organ long axis. Wrinkles in the epidermis run parallel to the long axes of epidermal cells and are absent outside of the immediate regions of pulvini.

The shapes of epidermal cells of primary pulvini transitioned from oblong and perpendicularly oriented in the region of the pulvinus to oblong and parallelly oriented in the region of the petiole (Fig. 2). Epidermal cells in the intermediate region between pulvinus and petiole displayed relatively circular aspect ratios (Fig. 2b) and did not display an obvious orientation relative to the organ long axis (Fig. 2c). The average epidermal cell aspect ratio in the region of the pulvinus was 3.70 ± 1.15 (mean ± SD) oriented at an average angle of 88.2° ± 6.85° (mean ± SD) relative to the organ long axis.

**Figure 2:**
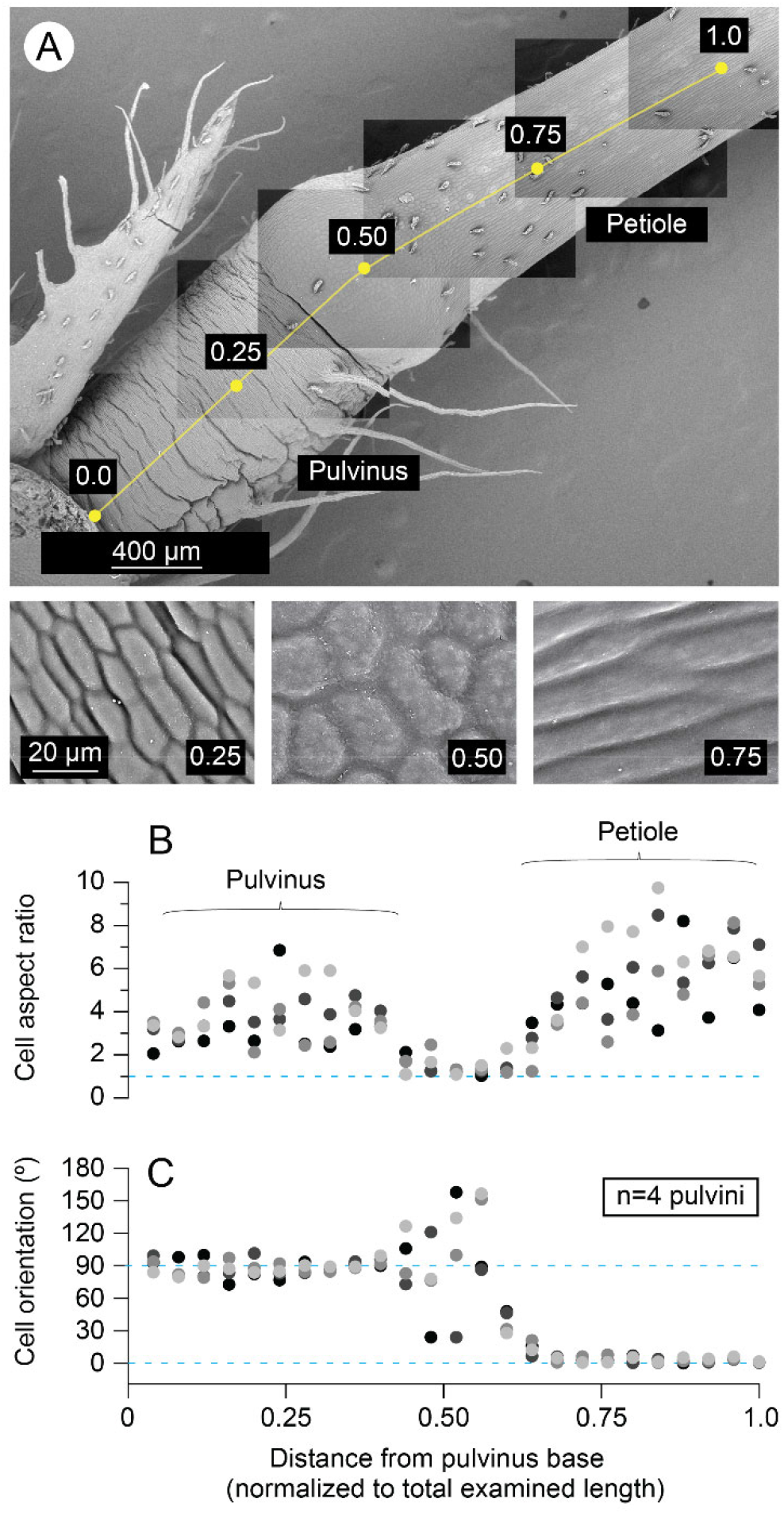
Spatial variation in epidermal cell morphology. [A] Stitched scanning electron micrographs showing a primary pulvinus organ in lateral view. Yellow line depicts a path traced along the central axis of pulvinus and petiole. Inset micrographs show epidermis morphology at higher magnification. Labels on inset micrographs indicate the approximate location at which each was taken, expressed as a portion of the total path length examined. [B] Epidermal cell aspect ratio varies across the lengths of pulvini and petioles. Dashed blue line denotes a circular aspect ratio of 1. [C] Epidermal cell orientations expressed relative to the orientation of the central path. Dashed blue lines fall at 90° and 0° denoting perpendicular and parallel cell orientations, respectively. Data points collected from the same pulvini are depicted in matching shades of gray in both B and C.

High magnification scanning electron microscopy allowed visualization of individual cellulose microfibrils at the inner surfaces of parenchymal cell walls (Fig. 3). Cellulose microfibrils were aligned primarily dorso-ventrally in the plane of the wall with a predominant microfibril orientation of 82.5° ± 29.1° (mean ± SD) relative to the orientation of nearby vasculature.

**Figure 3:**
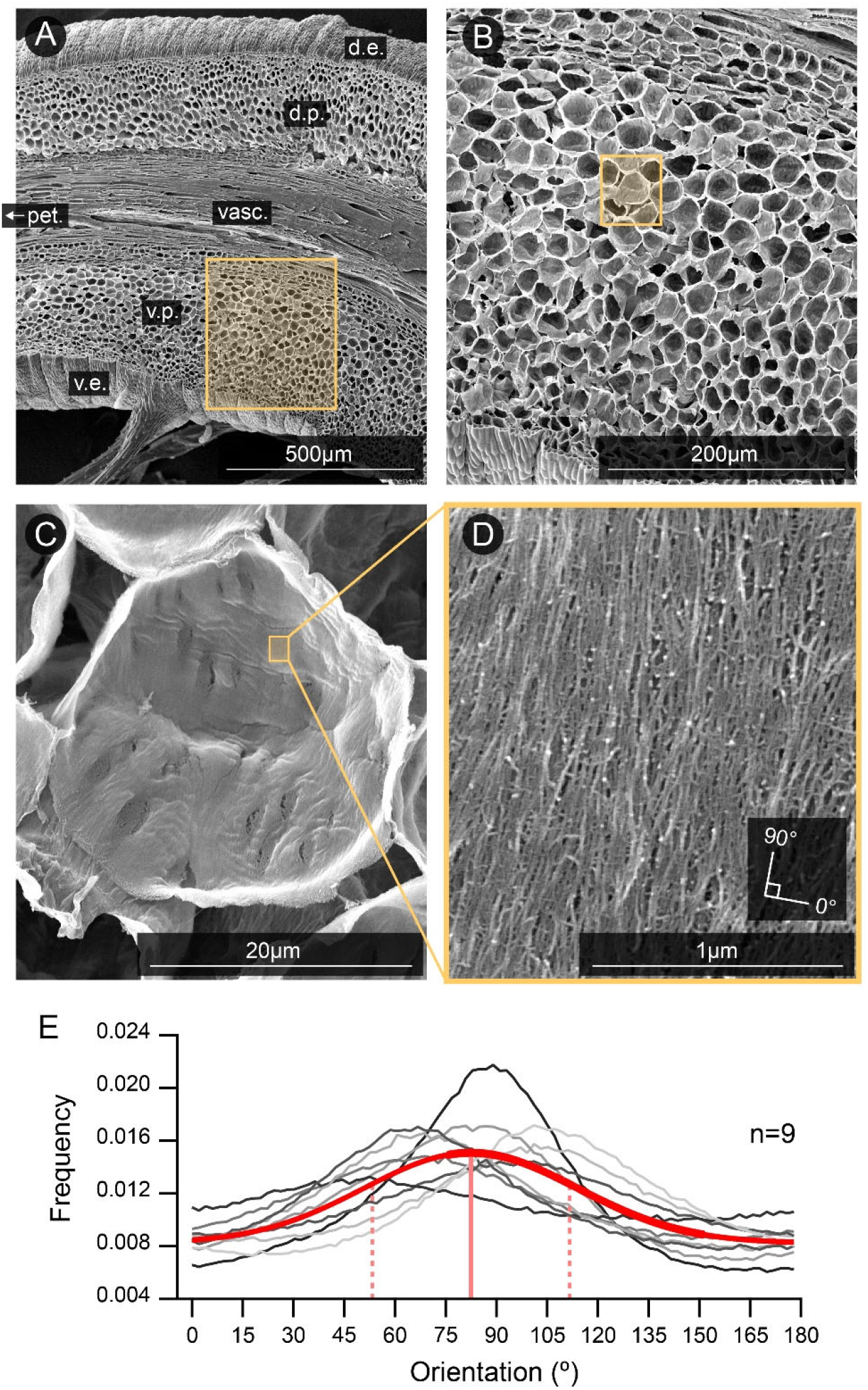
Orientations of cellulose microfibrils reinforcing parenchymal cell walls. [A] Scanning electron micrograph showing a lateral view of a longitudinally bisected primary pulvinus. Protoplasts have been removed chemically to reveal architecture at the inner face of the cell wall. Labels indicate dorsal, i.e. adaxial, epidermis (d.e.), dorsal parenchyma (d.p.), vasculature (vasc.), ventral, i.e. abaxial, parenchyma (v.p.), and ventral epidermis (v.e.). Organ is oriented with petiole (pet.; not visible) left of the frame. [B] Magnified view of the highlighted region in A showing ventral parenchyma bounded by vasculature and ventral epidermis. [C] Magnified view of the highlighted region in B showing the inner face of one parenchymal cell wall. [D] High magnification view of the highlighted region in C showing cellulose microfibrils running primarily dorso-ventrally in the plane of the cell wall. Fiber angles are described relative to the long axis of nearby vasculature (defined as 0°) with 90° indicating a perpendicular orientation. [E] Histograms showing distributions of fiber orientations in 9 cells from 4 pulvini. Each gray trace shows the distribution of fiber angles in a high magnification scanning electron micrograph similar to D. A gaussian curve (solid red trace) fit to all histograms indicates a predominant fiber orientation of 82.5 degrees (solid vertical line) with a standard deviation of ± 29.1 degrees (dotted vertical lines).

Consistent with the necessity for rapid fluid exchange during active movement, primary pit fields connecting adjacent parenchyma cells were prevalent in pulvini of *M. pudica*. Pit fields displayed a mix of circular, oval, and spindle-shaped morphologies (Fig. 4). In many parenchyma cells, pit fields were predominantly oval or spindle-shaped with long axes oriented perpendicular to the organ axis. For example, pit fields in the five parenchyma cells depicted in figure 4b displayed aspect ratios ranging from 1.2 to 4 with an average aspect ratio of 2.22 ± 0.78 (mean ± SD, n=30 pit fields) and an average long axis orientation of 93.0 ± 25.0 degrees (mean ± SD) relative to the organ axis. Parenchyma cells with predominantly oval or spindle-shaped pit fields were often directly adjacent to cells with predominantly circular pit fields, indicating that parenchyma cells of similar size, shape, and position can possess pit fields of varying morphology.

**Figure 4:**
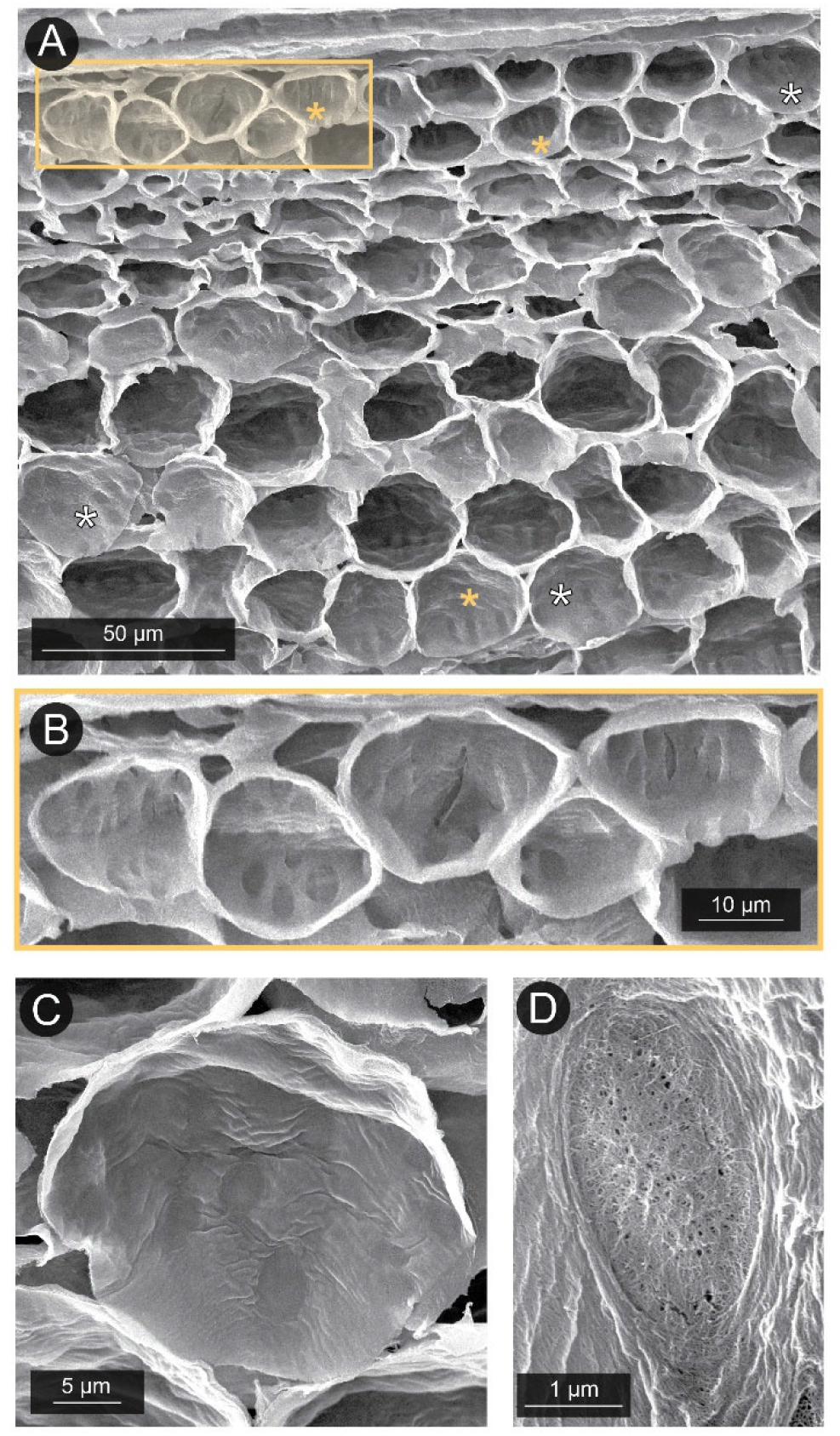
Primary pit fields visible in the walls of decellularized parenchyma cells. [A] Scanning electron micrograph showing a lateral view of a longitudinally bisected primary pulvinus. The organ long axis is oriented horizontally with the central vascular core visible at the top of the panel. Pit fields are visible as darkened regions of cell wall and display a mix of circular, oval, and spindle-shaped morphologies. Yellow asterisks highlight cells displaying predominantly oblong pit fields oriented perpendicular to the long axis of the pulvinus. White asterisks highlight cells with more circular pit field morphologies. [B] Magnified view of the highlighted region in A showing parenchyma cells with predominantly spindle-shaped pit fields oriented perpendicular to the organ long axis. [C] A single parenchyma cell displaying a mix of circular and oval shaped pit fields. [D] A single spindle-shaped pit field composed of mat-like cellulose microfibrils perforated by numerous plasmodesmata channels.

Osmotic swelling of dissected pulvinus organs and tissue samples yielded anisotropic deformation across all scales of pulvinus organization studied (Fig. 5). Across a graded series of osmotic baths ranging from 0 to 878 mOsm, cylindrical sections of pulvinus organ with an intact epidermis (Fig. 5c) displayed a 15.3 ± 2.75% (mean ± SD, p < 0.01) change in length without significant changes in either width (1.74 ± 1.83%, p = 0.11) or height (1.02 ± 2.42%, p= 0.10). Cylindrical pulvinus sections skinned of their epidermis (Fig. 5d) varied in length 17.7 ± 4.26% (p < 0.01), increased marginally in width (2.91 ± 1.98%, p < 0.01), and displayed increased variation in height (7.80 ± 8.55%, p = 0.05). The increased variation in the height of cylindrical organ sections skinned of their epidermis was attributable to a tendency for elongating parenchyma tissues to bow dorsally and ventrally away from the central vasculature of the pulvinus organ. This bowing resulted in skinned pulvinus sections adopting spindle-shaped lateral profiles reminiscent of rugby or American footballs when swollen osmotically (Fig. 5f). Isolated blocks of parenchyma tissue dissected free of epidermis and vasculature (Fig. 5e) changed in length 26.2 ± 3.15% (p < 0.01) without statistically significant changes in either width (−1.39 ± 2.40%, p = 0.26) or height (3.76 ± 5.75%, p = 0.22).

**Figure 5:**
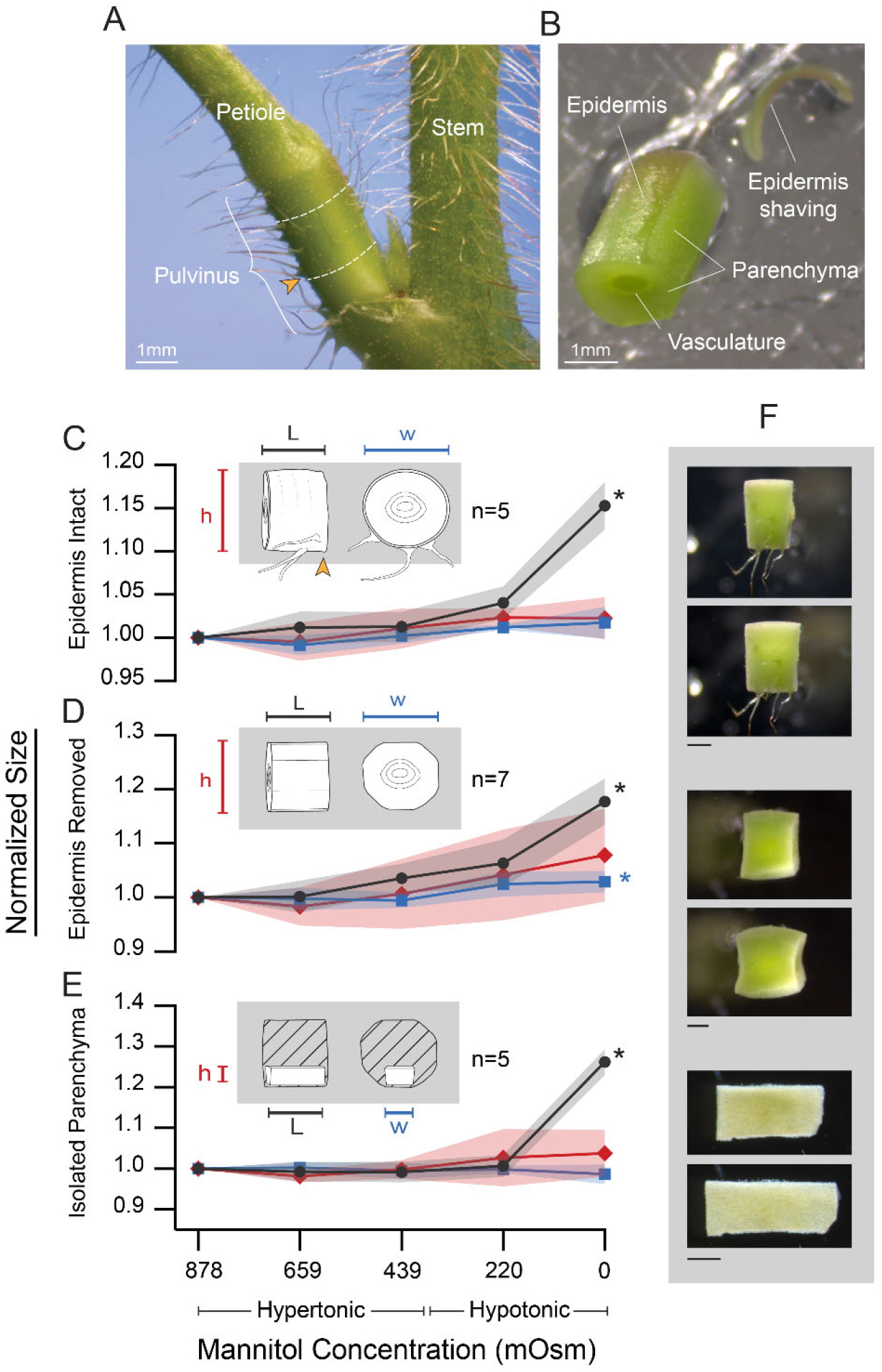
Osmotic perturbations reveal anisotropic swelling behaviors across scales of organization. [A] Lateral view of an intact *Mimosa pudica* primary pulvinus with flanking leaves removed. Dashed white lines illustrate cross-sectional cuts used to isolate a cylindrical section of pulvinus. Orange arrowheads in A and C indicate proximal, ventral corner of sections. [B] Isolated cylindrical section of pulvinus with epidermis partially removed to illustrate skinning technique. [C, D, E] Effect of graded osmotic bathing solutions on the length (L, black circles), cross-sectional width (w, blue squares), and height (h, red diamonds) of excised pulvinus sections. Inset line drawings depict dimensions and typical morphologies in lateral and cross-sectional views. Graph markers are means ± SD. Data is normalized to tissue dimensions in the most hypertonic bathing solution (878 mOsm). Asterisks denote values significantly different from 1.0 in the most hypotonic bathing solution (0 mOsm). [C] Isolated pulvinus sections expand longitudinally in increasingly hypotonic bathing solutions with negligible change in width or height. [D] Sections manually skinned of their epidermis expand primarily in length but display a small increase in width and increased variability in height. [E] Isolated blocks of parenchyma tissue expand longitudinally with negligible change in width or height. Sample sizes (n) for each condition are noted beside inset illustrations. [F] Representative images illustrating shape changes of pulvinus preparations. Upper and lower images of each pair display sample shape in hypertonic and hypotonic baths, respectively. Scale bars 500 μm.

The geometries of live parenchyma cells visualized by confocal microscopy depended on the tonicity of the solution in which they were bathed (Fig. 6). In isotonic bathing solution, the average ratio of cell width to cell height was 0.95 ± 0.15 (mean ± SD), which does not differ significantly from a circular cross-sectional aspect ratio of 1.0 (p=0.54). The average ratio of cell length to cell height was 0.85 ± 0.27 (mean ± SD), which differs significantly from a circular lateral aspect ratio of 1.0 (p=0.01), confirming that live parenchyma cells bathed in isotonic solution are approximately disk-shaped; circular in cross section but flattened longitudinally in the direction parallel to the long axis of the pulvinus. Cells reimaged in osmotically hypotonic distilled water baths did not change significantly in width (−0.56 ± 1.88%, mean ± SD, p = 0.13), displayed a small but significant change in height (1.78 ± 2.99%, p < 0.01), and displayed a relatively large increase in length of 14.5 ± 5.07% (p < 0.01) occurring along the long axis of the pulvinus (Fig. 6b). Following transfer to hypotonic bathing solution, cells displayed an average ratio of cell length to cell height of 0.96 ± 0.30, which was not significantly different from circular (p=0.54), indicating that longitudinally flattened, disk-shaped cells expanded towards more perfectly circular lateral profiles under increased turgor.

**Figure 6:**
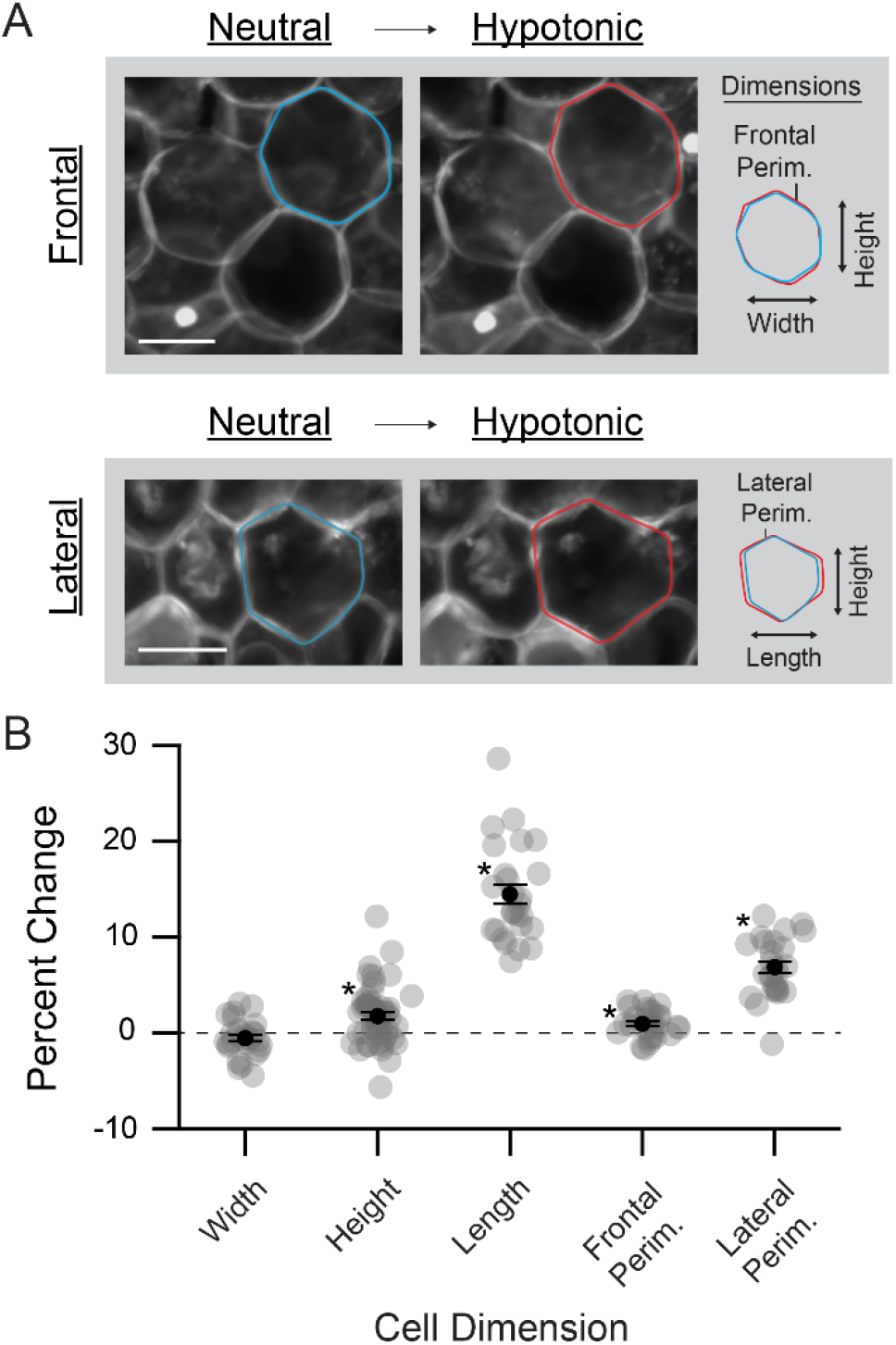
Nonuniform osmotic swelling of individual parenchyma cells. [A] Confocal micrographs of parenchyma tissue imaged in osmotically isotonic (384 mOsm) and hypotonic (distilled water) baths. Representative micrographs show cell morphology as seen in frontal and lateral views. All scale bars 20 μm. Inset line drawings illustrate anatomical dimensions and highlight changes in cell shape between osmotic baths. [B] Effect of osmotic swelling on frontal and lateral dimensions of 53 cells from 8 pulvini. Gray dots represent percent changes in cell dimensions following transfer from isotonic to hypotonic bath. Black markers represent means ± standard errors. Asterisks denote percent changes significantly different from zero (p<0.05).

Measurements of frontal and lateral cell perimeters revealed anisotropy in the amount of cell wall distension that occurred during osmotic expansion. Lateral cell perimeter, measured in the plane parallel to the organ long axis, increased 6.87 ± 3.10% (p < 0.01) following acclimation in hypotonic bathing solution. In contrast, frontal cell perimeter measured in cross section perpendicular to the organ long axis changed only marginally (0.96 ± 1.41%, p < 0.01). This difference in cell wall distension under isotropic turgor rise indicates that the parenchyma cell wall is materially anisotropic, displaying greater extensibility in the longitudinal direction than in the circumferential.

## Discussion

The hydraulic nature of pulvinus-driven movement has been recognized since the 1800s and is discussed in Darwin’s *The Power of Movement in Plants* (Darwin and Darwin, 1880). However, an understanding of the functional morphologies that transform isotropic cellular turgor pressures into anisotropic, directed movement at the organ scale has remained fragmented. Here we demonstrate that in pulvini of *Mimosa pudica*, multiple anisotropic structural features are present across subcellular, cellular, and tissue levels of organization. These include perpendicularly oriented cellulose microfibrils and primary pit fields in parenchymal cell walls, longitudinally flattened parenchyma cell geometries, and perpendicularly oriented epidermal cells associated with transverse surface wrinkles. We observed highly anisotropic osmotic swelling of parenchyma cells and tissues; however, swelling behaviors at higher levels of organization were also influenced by the presence of the enveloping epidermis, suggesting that hydraulic deformation of pulvini is influenced by structural anisotropy at multiple hierarchical scales.

### Cell wall anisotropy

Consistent with polarized light investigations of pulvinus morphology in the scarlet runner bean *Phaseolus coccineus L* (Mayer et al., 1985), scanning electron micrographs of decellularized *M. pudica* pulvini revealed cellulose microfibrils oriented predominantly perpendicular to the long axes of pulvini. Cellulose microfibrils are crystalline structures composed of closely packed chains of parallelly-oriented cellulose molecules (Brett, 2000) and display high tensile strengths and longitudinal stiffnesses of approximately 1 GPa and 130 GPa respectively (Gibson, 2012). Cellulose microfibrils function as the primary mechanical component of the plant cell wall, ultimately providing the tensile strength needed to balance large intracellular turgor pressures ranging from 0.4-0.8 MPa in typical hydrated plant tissues (Taiz et al., 2015) to over 1 MPa in some specialized plant cells (Franks, 2003).

Anisotropic patterns of cellulose reinforcement are not unique to pulvini. Cellulose orientation is actively controlled by the microtubule-guided motion of cellulose synthase complexes that extrude cellulose microfibrils into the extracellular space (Brett, 2000), and anisotropic cellulose deposition is the primary means by which mechanical anisotropy is produced and maintained in plant cell walls. The interplay of intracellular turgor pressure and cell wall anisotropy ultimately governs the direction of irreversible turgor induced shape changes (Baskin, 2005) and underlies the development of complex non-spherical morphologies (Bidhendi and Geitmann, 2016) in growing plant cells. Interactions between cell wall anisotropy and turgor pressure have also been shown to underlie reversible deformation of certain mature plant cell types. The hydraulic guard cells flanking stomatal pores, for example, are reinforced by cellulose microfibrils oriented primarily perpendicular to the guard cell long axes (Galatis and Apostolakos, 2004). The perpendicular orientation of microfibrils inhibits radial expansion of guard cells while permitting elongation during hydraulic expansion, ultimately facilitating the microscopic outward bowing of guard cells that facilitates repeated opening and closing of stomatal pores (Aylor et al., 1973). Parenchyma cells observed in the current study displayed a high degree of directional anisotropy in the extensibilities of their cell walls, with osmotic expansion causing wall distension approximately 7 times greater in the longitudinal direction than in the circumferential. Analogous to the effect of cellulose anisotropy in stomatal guard cells, this finding suggests that cellulose microfibrils selectively restrict circumferential expansion of parenchymal cell walls in the pulvini of *M. pudica*, resulting in predominantly longitudinal cell wall deformation in response to turgor change.

In addition to unidirectionally oriented cellulose microfibrils, frequent primary pit fields also have the potential to contribute to mechanical anisotropy in the parenchyma cell wall. Primary pit fields are typically circular, sometimes oval regions of the primary cell wall perforated by numerous plasmodesmata. In addition to being perforated, the cell wall is significantly thinner in pit fields compared to the surrounding wall, commonly by a factor of 5 or more (e.g. Kobiyama and Crandall-Stotler, 1999). The presence of numerous primary pit fields is consistent with the importance of rapid water exchange between cells in the *M. pudica* pulvinus; however, the presence of many oblong and spindle-shaped pit fields oriented perpendicular to the organ axis suggests their shape and orientation may also serve a mechanical function. Assuming the reduced thickness of the pit field is associated with a lower elastic modulus, oblong pit fields oriented perpendicular to the pulvinus axis may serve as expandable seams that facilitate distension of the cell wall preferentially in the longitudinal direction. Direct measurements of pit field deformation or elastic modulus are needed to verify this hypothesis, and exploration of the influence of pit field material properties and shape on cell wall mechanics may prove a fruitful topic for future research.

### Geometric deformation of parenchyma cells

While perpendicularly oriented cellulose microfibrils and pit fields are conducive to long-axis deformation of parenchyma cells, our results indicate that direct stretching of the cell wall cannot account for all of the longitudinal cell deformation observed in the current study. Following transfer from isotonic to hypotonic bathing solution, the average length of parenchyma cells increased by 14.5 ± 5.07% (mean ± SD); however, the lateral perimeters of these same parenchyma cells increased by only 6.87 ± 3.10%, indicating that direct stretching of the cell wall accounts for less than half of longitudinal cell deformation on average, even under the assumption that all cell wall distension contributes exclusively to elongation. Geometric deformation of cells from longitudinally flattened lateral profiles towards more longitudinally expanded lateral profiles likely accounts for the remaining observed cell deformation. Like the corrugated bellows of an accordion or concertina, the longitudinally flattened morphologies of parenchyma cells are conducive to longitudinal expansion and contraction via geometric changes in cell shape. We observed both direct distension of the cell wall and geometric shape changes of cells under increased turgor, indicating that both of these mechanisms contributed to anisotropic parenchyma cell deformation during experimentally imposed swelling. Further studies of parenchyma cell deformation at physiologically realistic intracellular turgor pressures are needed to determine the relative contributions of these two mechanisms to anisotropic deformation *in vivo*.

Here we focus exclusively on parenchyma cells from the relatively large extensor (i.e. ventral or abaxial) cortex of the primary pulvinus of *M. pudica*. Comparison with morphology and swelling behavior of the flexor cortex was not carried out in the current study but is a worthwhile topic for future research. We expect that parenchyma cells of the flexor cortex would display similar structural anisotropy and directional swelling behavior to that of the extensor cortex; however, subtle differences in morphology across cortexes likely influence their behaviors. In pulvini of the scarlet runner bean *Phaseolus coccineus L*., for example, both extensor and flexor cortexes are composed of longitudinally flattened, disk-like parenchyma cells. Cells of the extensor compartment are notably larger in size, and isolated blocks of extensor parenchyma tissue display relatively larger percent changes in length than flexor tissues when osmotically swollen in distilled water (Mayer et al. 1985). Such differences may be relevant to pulvinus mechanics *in vivo*, and further investigation may provide insight into the granularity of morphological specialization displayed by parenchyma cells that are subject to differing functional demands.

### Mechanical contribution of the pulvinus epidermis

Most plant stems and petioles are ensheathed in a skin of elongate epidermal cells oriented parallel to the long axis of the organ (Kutschera, 2008). This parallel orientation of epidermal cells is a natural consequence of turgor-driven morphogenesis, in which elongated plant organs form developmentally through anisotropic expansion and elongation of the cells that compose them. Over most of their lengths, the petioles of *M. pudica* display a typical epidermis morphology of elongate cells oriented parallel to the organ long axis. Surprisingly, however, we observed epidermal cells oriented perpendicular to the growth axis in the immediate regions of pulvini. Perpendicular cells correlated spatially with the presence of transverse wrinkles oriented perpendicular to the long axis of the pulvinus. Transverse wrinkles in the epidermis of *Mimosa pudica* have been described previously and have been shown to expand and contract dynamically during active movement of live plants (Song et al., 2014). They are thought to provide extra surface area that increases the longitudinal extensibility of the epidermis without compromising its circumferential stiffness (Mano and Hasebe, 2021).

We suggest that perpendicularly oriented epidermal cells are a morphological specialization that facilitates transverse wrinkling of the epidermis by reducing its effective flexural stiffness in the longitudinal direction. Epidermal cell orientation has been shown to have a direct effect on the material stiffness of plant epidermis, inducing anisotropic tensile properties and relatively lower tensile stiffness in the direction perpendicular to the orientation of epidermal cells (Bidhendi et al. 2020). Importantly, a perpendicular orientation of epidermal cells also allows transverse wrinkles to form between epidermal cells rather than across their lengths, minimizing bending strains imposed on individual epidermal cells. Direct mechanical tests exploring the impact of epidermal cell orientation on flexural stiffness are needed; however, facilitation of transverse wrinkle formation provides a potential explanation for the observation of perpendicularly oriented epidermal cells confined to the immediate regions of pulvini.

In cylindrical sections of pulvinus organ skinned of an enveloping epidermis, we observed outward bowing of parenchyma tissue away from the central axis of cylindrical pulvinus sections during osmotic swelling. We suspect that this parenchymal bowing results from a difference in the longitudinal extensibility of parenchyma tissues and the vascular tissue they surround. Isolated parenchyma tissues dissected away from the mechanical context of the pulvinus organ display a strong bias for purely longitudinal expansion; however, when coupled to a relatively stiff vasculature core in the context of an intact organ, pure longitudinal expansion of the parenchyma along a straight line is resisted, and outward curvature of longitudinally expanding parenchyma tissues results. We did not observe the phenomenon of outward parenchyma bowing in pulvinus preparations with an intact epidermis, suggesting that the circumferential stiffness of the epidermis resists outward bowing of parenchyma tissues. While anisotropic cell wall reinforcement and cell geometry are well suited to ensure longitudinal expansion at the scale of individual parenchyma cells, our findings suggest that the transversely wrinkled epidermis serves as an encompassing sleeve that restricts radial deformations that emerge at higher scales of tissue organization, ensuring that hydraulic expansion of parenchyma tissue *en masse* remains aligned along the long axis of the pulvinus where its effect on organ bending is greatest.

### Osmotic swelling as a model for *in vivo* physiology

Simple osmotic swelling and shrinkage of pulvinus tissues does not replicate the rapid and complex mechanisms of intercellular fluid movement employed by *Mimosa pudica* pulvini *in vivo*, the details of which are not fully understood. Rather, osmotic perturbations employed in the current study provide a means of introducing purely isotropic, nondirectional intracellular loads into parenchyma cells in the form of hydrostatic fluid pressures, the effects of which provide insight into multiscale tissue mechanics. While the precise physiological mechanisms powering active fluid movement and turgor change *in vivo* is an important topic for further investigation, here we show that hydraulic deformation of *M. pudica* pulvini is a product not only of biochemical specialization, but also of a suite of downstream structural features that span multiple hierarchical scales of organization. These features are ideally situated to translate isotropic, non-directional cellular turgor pressures into the physiologically relevant motor task of long-axis organ bending, suggesting that they are important contributors to the mechanism of pulvinus-driven motion *in vivo*.

## Supporting information

Supplementary Material Titles and Captions

## Acknowledgements

We thank Youssef Chebli for help in microscope operation and helpful discussions about data collection, and David Liu and S. Kelly Sears at the Facility for Electron Microscopy Research of McGill University for help in microscope operation and data collection.

## Data availability

Tabulated data from all figures are available as supplemental files.

