## Supplementary Material Titles and Captions for "Multiscale structural anisotropy in pulvinus motor organs of the sensitive plant *Mimosa pudica*"

Titles and captions of supplemental files for “Multiscale structural anisotropy in pulvinus motor organs of the sensitive plant *Mimosa pudica*”

Supplemental Datasets

**Table S1:** Epidermis cell aspect ratios and orientations

Caption: Tabulated values used to build figure 2, panels B and C. Cell position is expressed as a portion of the total organ length examined. Cell orientation is expressed relative to the long axis of pulvinus and petiole.

**Table S2:** Cellulose microfibril orientation histogram data

Caption: Tabulated output of the Directionality plugin of ImageJ, FIJI. These data were used to build the histograms in figure 3, panel E. All angles are expressed relative to the orientation of the central vascular bundle, which is defined as 0 degrees. Angles increase in a counterclockwise direction. Tabulated angles are lower bounds of histogram bins.

**Table S3:** Pit field dimensions and orientations

Caption: Tabulated measurements of pit fields from parenchyma cells visible in figure 4, panel B. Pit field orientation is expressed relative to the orientation of nearby vasculature.

**Table S4:** Dimensions of dissected pulvinus preparations

Caption: Tabulated dimensions of dissected pulvinus preparations across a graded series of osmotic baths. These data were used to build figure 5, panels C, D, and E.

**Table S5:** Parenchyma cell dimensions

Caption: Tabulated dimensions of individual parenchyma cells in osmotically isotonic and osmotically hypotonic baths. These data were used to build figure 6, panel B.
